# Characterization of real B_0_ shim fields ^generated^ by higher order B_0_ shim systems of whole body human 3T and 7T MRI systems

**DOI:** 10.1101/2025.08.26.672172

**Authors:** Mahrshi Jani, Shengyue Su, Lena Gast, Armin Nagel, Anke Henning

## Abstract

**Purpose:** Ensuring optimal homogeneity of the static magnetic field (B0) is critically important for significantly enhancing the image quality, spectral resolution, and overall diagnostic accuracy of magnetic resonance imaging (MRI) and magnetic resonance spectroscopy/spectroscopic imaging (MRS/MRSI), especially at high and ultra-high field strengths. Achieving this goal relies on the deployment of advanced, higher-order shim hardware, which is indispensable for effective and precise B0 shimming.

**Methods:** In this study, we acquired B0 field maps to characterize the performance of each shim channel of vendor-provided B0 shim hardware integrated into whole body human 7T MRI system. To rigorously evaluate and compare the B0 field homogeneity achievable using these devices, we utilized spherical harmonic expansions to fit the measured B0 distributions for each shim channel in each of the investigated built-in B0 shim systems. This methodology enabled a systematic assessment of shim field “purity” and the effectiveness of different B0 shim configurations.

**Results:** While our analysis confirmed that the linear-order shim terms provided by the gradient coils maintained high purity, certain second-order shim terms and all third-order B0 shim coils of all vendors 7T whole body human MRI systems exhibit a high level of impurity and more specifically introduce substantial linear field components. These unintended linear contributions interfere with the linear shim fields provided by the gradient coils negating their corrective influence. As a result, the 3_rd_ order B0 shim coils and some 2^nd^ order B0 shim coil elements compromised the overall effectiveness of the B0 shimming process rather than improving it. Although pure higher order B0 shim fields were previously shown to substantially improve B0 field inhomogeneity, the vendor implemented versions of higher order B0 shim systems in current 7T whole-body human MRI scanners cannot achieve the desired level of uniformity and partly perform worse than first order B0 shimming. Substantial deviations between 2^nd^ order and especially 3^rd^ order real shim fields versus ideal shim fields demonstrate the necessity of real shim field calibration and call for improvement of future 2^nd^ and 3^rd^ order B0 shim coil design for human whole-body MRI scanners. Our findings offer valuable guidance for optimizing B0 shimming strategies at high and ultra-high field strengths, ultimately enhancing image quality.

**Conclusion:** Recent evaluations have demonstrated that certain second- and third-order shim coils, as implemented in current commercial 7T human whole-body MRI systems, introduce unintended lower-order (linear) field components. These field “impurities” substantially deviate from the desired higher-order spatial harmonics, thereby diminishing the orthogonality of the higher order B_0_ shim system. This in turn largely compromises the effectiveness of higher-order B_0_ shimming at whole-body 7T MRI scanners. These findings emphasize the necessity of conducting accurate shim field calibration to best utilize the current 7T B0 shim system configurations on one hand. On the other hand, improving the purity of these commercial B_0_ shim systems in the design phase to ensure that the higher-order terms accurately match their intended spherical harmonic profiles, particularly for the third-order shim terms—would largely enhance the achievable B_0_ field homogeneity at 7T. This improvement has directly clinically significant outcomes.

## INTRODUCTION

The shift towards ultra-high field strengths of 7 Tesla and above human whole-body MRI systems offer significant benefits for various clinical and research applications including functional MRI (fMRI) and magnetic resonance spectroscopy (MRS) due to increased signal-to-noise ratio, spectral separation and enhanced blood oxygen level dependent (BOLD) effect. Nevertheless, in order to leverage these benefits, it is imperative for the static magnetic field (B_0_) to exhibit high homogeneity^1,2^. When an object or human subject is introduced into the magnetic field, variations in B_0_ field homogeneity arise, primarily caused by variations in magnetic susceptibility among different types of tissue and at interfaces between tissue and air ^3^.

The homogeneity of the B_0_ field plays a critical role in magnetic resonance imaging and spectroscopy with B_0_ inhomogeneity leading to various aberrations including distortion in shape, reduction in image quality, signal loss, and deterioration in spatial resolution in anatomical and functional MRI. Echo planar imaging (EPI) sequences exhibit particular vulnerability to various types of these imaging distortions ^4^. In addition, challenges such as broadening of spectral lines, localization errors, decline in spectral resolution and failure of water and fat suppression techniques will result from B_0_ inhomogeneity in spectroscopy experiments ^5,6,7^.

Current 7T state-of-the-art B_0_ shimming methods either extend spherical harmonic (SH) terms to third ^8^- or even fourth-degree fields^9^ or adopt localized multi-coil shim arrays^10^. While multi-coil systems offer more degrees of freedom due to their numerous small coils (e.g., 48 coils in Juchem et al.^11^), SH-based B_0_ shimming benefits from readily available, orthogonal B_0_ shim fields that follow well-defined analytical models. These SH systems are commonly integrated into standard MRI hardware or can be obtained commercially. The acquisition procedures and shim algorithms for B_0_ shimming^12–20^have been thoroughly examined in numerous previous studies. Both multi-coil and spherical harmonic B_0_ shimming strategies require accurate characterization of the actual B_0_ shim fields produced by each shim coil^12–15^. In the multi-coil approach, a large array of small coils provides substantial flexibility in shaping the B_0_ field due to the increased number of adjustable coil currents. In contrast, spherical harmonic B_0_ shimming typically leverages hardware that is readily integrated into the MRI scanner or available commercially ^21^ While some 1.5 T and most 3 T human MRI scanners are typically equipped with second order SH B_0_ shim coils, the majority of 7T human MRI scanners incorporate 2^nd^ and 3^rd^ order SH shim coils. The underlying SH shim coil designs are based on analytically defined spherical harmonic expansions (derived from Legendre polynomials), which produce inherently orthogonal B_0_ shim fields. This property facilitates vendor-supplied optimization algorithms to determine the best possible shim current settings ^22,23^. Especially respective B_0_ shim algorithm implementations of human MRI scanner vendors assume that each B_0_ shim coil produces ideal spherical harmonic shim fields. However, if B_0_ shim fields are not ideal spherical harmonic functions, B_0_ shimming either needs to be done iteratively until sufficient convergence is achieved ^24^ or alternatively, reference field maps can be acquired for each B_0_ shim coil to characterize and consider non-ideal shim field contributions already during the computation of shim currents ^17, 20^. This real shim field calibration approach for SH shim systems has been successfully used in preclinical MRI systems ^18,25^ and with an insert shim system for human ultra-high field MRI systems to shim up to 4^th^ plus degree shim terms ^21,26^.

Beyond SH shimming Previous studies have also modeled real shim fields for B_0_ shimming using matrix shim coils ^16,27^. Furthermore, the impact of third-order SH shim coils on gradient field linearity and their interactions with gradient coils at ultra-high field human MRI systems (7 T and above) has been explored in earlier work^19^. Prior discussions have also touched upon the eddy current effect in different SH shim coils and provided a pre-emphasis calibration for dynamic SH shimming^18^. In addition, researchers have investigated the gradient impulse response function (GIRF) and corresponding shim impulse response functions through frequency sweep experiments^27, 28^. These efforts utilized advanced field monitoring devices, such as field cameras, to accurately capture and characterize the time-dependent properties of both gradients and shims^18,27, 28^. Stronger gradients and higher magnetic fields intensify their interaction and increase the equipment’s workload ^27,29^.

However, the literature currently offers limited insights into the shim field impurities introduced by standard vendor-supplied higher-order SH B_0_ shimming hardware at clinical 3T and 7T human whole-body MRI scanners. A rigorous characterization of these higher-order B_0_ shim fields in commercial clinical MRI systems and their influence on the MRI and MRS data quality remains largely unexplored.

In this study, we provide a qualitative and quantitative evaluation of the actual B_0_ shim fields generated by vendor-sourced SH B_0_ shim hardware operating under various 3T and 7T magnet configurations. We then compare the resulting shim field impurities from these in-bore SH B_0_ shim systems to those obtained using specialized “external” insert shim hardware ^21^, thereby offering new insights into the spatial fidelity and performance limitations of current vendor integrated B_0_ shimming technologies.

## METHODS

### Data Acquisition

This study utilized four human whole body MRI scanners from two different manufacturers (Siemens, Philips) with two different field strengths 3T and 7T, to investigate the purity of commercial SH B_0_ shim systems. For a field strength of 7T, three first order, five second order, and either seven (Philips) or four (Siemens) third order B_0_ shim coils were provided by the vendors. For the 3T system, both vendors have three first order and five second order B_0_ shim coils. The sensitivities of each shim channel from all four human MRI scanners can be found in Table 1.

For each B_0_ shim coil, reference field maps were acquired using a spherical silicon oil phantom with a 170-mm diameter, chosen to approximate the volume of an average human brain ^17,18^. The real B₀ shim field was characterized by acquiring B₀-maps on four different MRI scanners using 3D gradient-echo protocols. In all cases, the echo-spacing (ΔTE) was chosen to maintain fat–water in-phase coherence and to minimize phase-wrapping artifacts in the field map. The sequence parameters were as follows: **7 Tesla systems: Siemens 7T** : TE₁ = 1.00 ms, TE₂ = 3.06 ms (ΔTE = 2.06 ms), TR = 4.3 ms; flip angle = 3°; Voxel size = 4.4 × 4.4 × 4.4 mm³; 52 slices, Field of view (FOV) = 282 × 282 × 274 mm³. **Philips 7T :** Multi-acquisition (dual echo) with ΔTE = 1.0 ms; TR = 10 ms; flip angle = 20°; Voxel size = 3 × 3 × 3 mm³; 50 slices, FOV = 200 × 200 × 100 mm³. **3 Tesla systems, Siemens 3T:** TE₁ = 4.50 ms, TE₂ = 6.96 ms (ΔTE = 2.46 ms); TR = 651 ms; flip angle = 20°; Voxel size = 3 × 3 × 3 mm³; 60 slices FOV = 192 × 192 × 100 mm³. **Philips 3T:** Multi-acquisition (dual echo) with ΔTE = 2.3 ms; TR = 24 ms; flip angle = 20°Voxel size = 3 × 3 × 3 mm³; 50 slices; FOV = 200 × 200 × 150 mm³.

Across all systems the field map was computed from the phase difference of the two echoes, ensuring consistent fat–water phasing and robust suppression of phase wrapping.

Prior to further analysis, any necessary phase unwrapping was performed using FSL bet ^30,31,32^. To establish the relationship between the input current of the shim amplifiers and the resultant shim field strength for each coil, additional reference fields were acquired at multiple current amplitudes. This allowed for a quantitative determination of how field intensity scales with input current. Calibration of the B₀ shim coil array was carried out using five discrete current amplitudes, explicitly omitting those corresponding to the vendor-provided shim solution. For first order (linear) gradient coils, current settings were confined to within ±15 % of each coil’s full operational range, whereas second- and third-order shim coils were calibrated over approximately ±35 % of their dynamic range. Acquisition parameters and processing algorithms were optimized to suppress phase-wrapping artifacts in the resulting B₀ field maps. Sampling this full dynamic range allowed us to characterize and subsequently correct amplitude-dependent nonlinearities. Each acquired field map was subsequently fitted to spherical harmonic expansions extending from the first to the sixth order ^21,26^. These fits formed the basis for constructing comprehensive shim calibration matrices that describe the real shim field impurities, thereby enabling precise higher-order shimming and improved B_0_ homogeneity for brain imaging applications.

### Modeling Reference B_0_ Shim fields

As described in the methodology, reference maps to determine the actual shim fields were obtained using a small spherical phantom with dimensions akin to the human brain. This choice ensured that field modeling remained confined to the target volume of interest (VOI), thereby minimizing the influence of large-scale field inhomogeneity outside the VOI on the accuracy of the model. Additionally, reference fields for all B_0_ shim coils were generated at multiple input current amplitudes, enabling the detection and correction of potential nonlinearities in shim field amplitude response.

For each B_0_ shim coil, reference fields obtained at various input current levels were characterized using a spherical harmonic decomposition^18,29,30^. In this framework, the three-dimensional scalar field distribution, B_0_(r), can be expressed as a linear combination of three-dimensional spatial basis functions, F_n_(r), with r = (x, y, z) representing the position vector. Since the in-bore B_0_ shim coil designs inherently rely on spherical harmonics, the spherical harmonic functions F_n_ (r) naturally serveas the appropriate basis functions for modeling the shim fields ^21^.

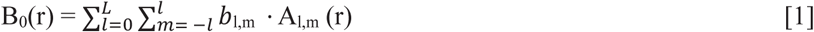

In this formulation, b_l,m_ are constants, where l denotes the spherical harmonic degree and m specifies the spherical harmonic order. The maximum degree of approximation, L, must be judiciously selected. While incorporating higher-degree spherical harmonic functions can refine the model, the gains eventually diminish, and beyond a certain point the model may begin to overfit the data. In this study, each measured B_0_ shim field was systematically modeled using spherical harmonic expansions ranging from the second through the sixth degree.

The spherical harmonic decomposition was performed by solving the standard over defined linear system of equations for n positions (in this case, up to sixth-degree spherical harmonic functions) ^21^:

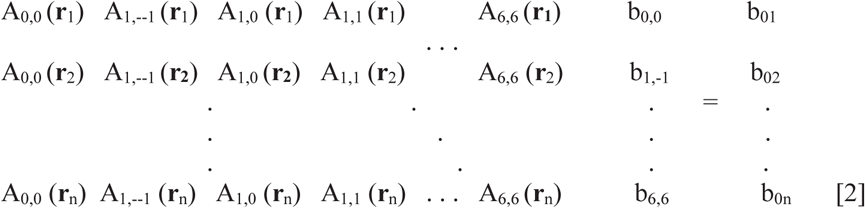

A_l,m_ is given by the Legendre polynomials. The values b_0k_ are the B_0_ values at the position **r**_k_, calculated from the B_0_ map:

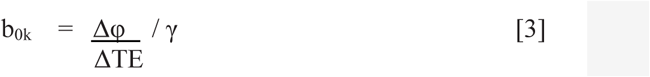

where γ is the gyromagnetic ratio, Δφ is the measured phase difference obtained from the acquired B_0_ maps, and ΔTE is the difference between the echo times of the two acquisitions of a B_0_ mapping sequence. Thus, the fields are modeled using a vector of coefficients b calculated from the linear system:

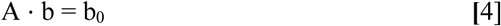

The B_0_ value at an arbitrary position r_k_ can be estimated by forming the A matrix and multiplying by the coefficient vector b.

This real B_0_ shim field modelling enabled the compilation of scanner specific B_0_ shim calibration matrices.

Real B_0_ shim fields modelled as described above were finally compared against ideal field distributions derived from the pure spherical harmonic functions of the same degree and order that the respective B_0_ shim coil was originally designed for. Respective difference maps have been computed.

Scatter plots were generated to visualize the relationship between the “real” shim field derived from the acquired shim field maps for each of the B_0_ shim coils (x) and it’s corresponding modeled “ideal” shim field using spherical harmonic functions (y). A least-squares linear regression line was fitted to each respective data set (y=β_0_+β_1_x), and the coefficient of determination (R^2^) was calculated as:

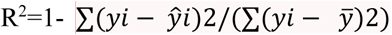

The R^2^ value, which denotes the proportion of variance between real versus ideal B_0_ shim fields of each investigated B_0_ shim coil was overlaid in the upper left corner of each plot alongside the regression equation to facilitate rapid visual assessment of the fit quality.

In addition, we performed a linear regression analysis to test the relationship between applied shim current and obtained real shim field amplitude for all investigated shim systems and shim coils.

To demonstrate the negative impact of shim field impurities of in-bore higher order SH shim coils we have finally computed B_0_ field maps after 1^st^ order, 2^nd^ order and 3^rd^ order B_0_ shimming of the whole brain assuming both vendors 7T real B_0_ shim fields, while applying a shim algorithm that assumes ideal fields just as the vendors do. In this context the maximum shim fields achieved with both systems is also reported and used in these simulations.

## Results

In comparing the measured B_0_ shim fields of each individual B_0_ shim coil of each vendors 3T and 7T human whole-body MRI system to their ideal spherical harmonic models, several key observations emerge regarding the performance and purity of the in-bore second- and third-order B_0_ shim coils from different vendors. Figures 1 and 2 illustrate these differences in the case of the first vendors (Philips Healthcare) 7T human MRI system (7T dSync). Here, the second-order B_0_ shim coil fields closely approximate their ideal spherical harmonic counterparts, exhibiting minimal deviations. Such fidelity ensures that the applied second-order B_0_ shimming is highly effective at reducing inhomogeneity in the B_0_ field. However, the examination of the first vendors third-order B_0_ shim coils reveal substantial linear field components that effectively introduce unwanted linear gradients. These strong linear impurities of the third-order B_0_ shim coils act in opposition to the linear B_0_ shims set by the gradient coils thus counteracting the beneficial effects that higher-order B_0_ shimming is supposed to achieve. Specifically, the linear contamination can diminish the net gain in field homogeneity that is anticipated from introducing these higher-order shim terms.

**Figure 1.**
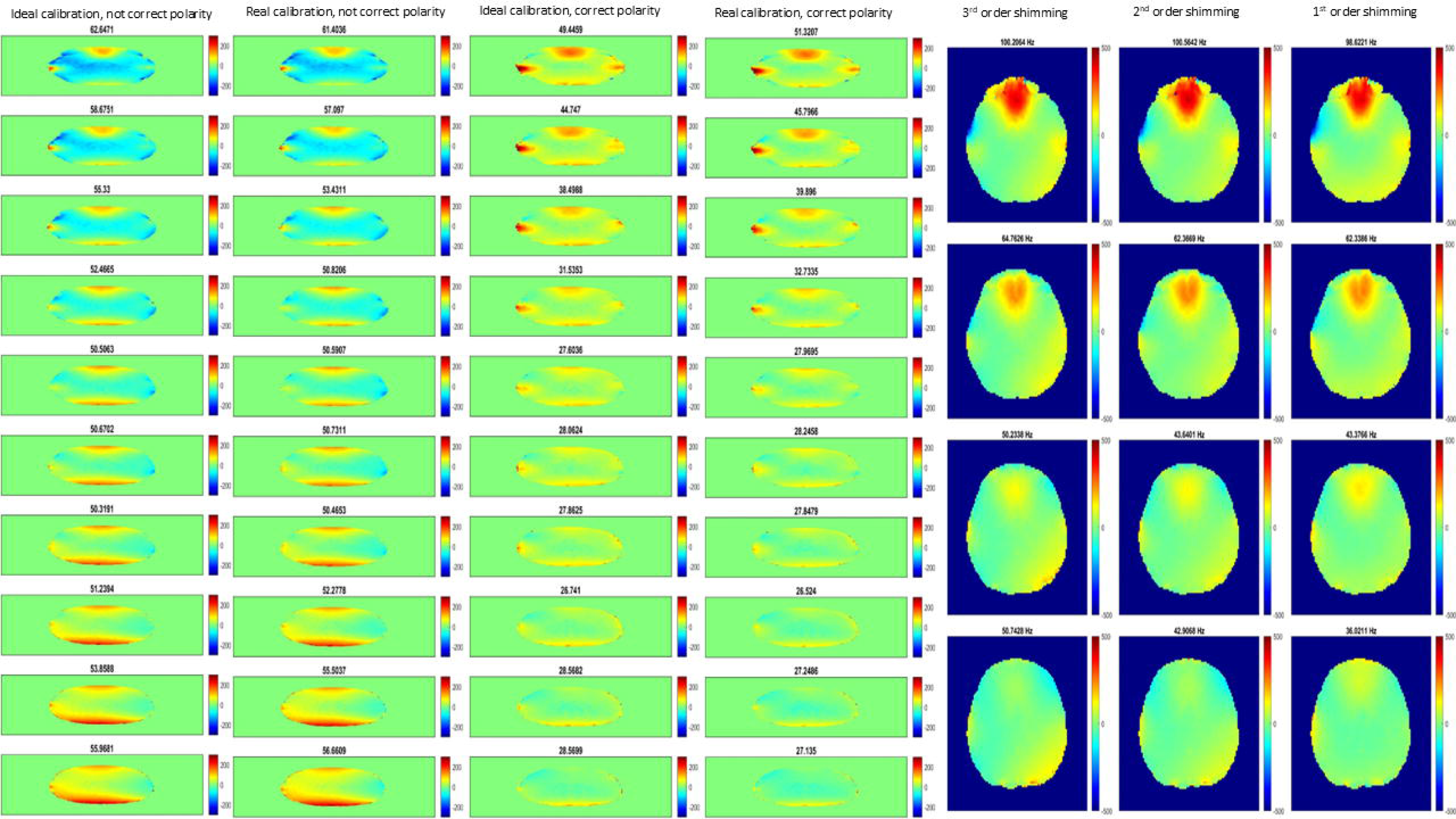
shows the real B_0_ shim fields (top row) for the first and second order SH B_0_ shim coils from the 7T Philips dSync MRI system (vendor 1), the respective ideal SH fields (middle row) and the difference maps (bottom row). Most of the real shim fields are almost identical to the respective ideal shim field models.

**Figure 2.**
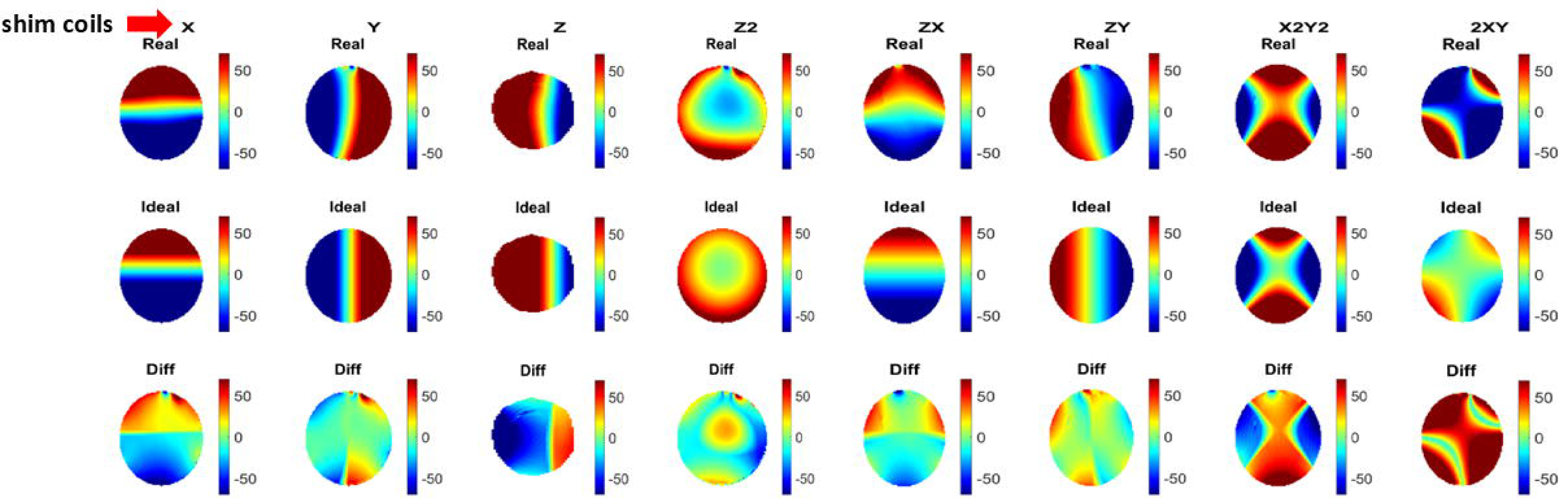
shows the real B_0_ shim fields (top row) for the third order SH B_0_ shim coils from the 7T Philips dSync MRI system (vendor 1), the respective ideal SH fields (middle row) and the difference maps (bottom row). Substantial linear field components are present in the third-order real shim coils and the real third order shim fields have hardly any resemblance with the theoretical SH model.

Figures 3 and 4 present the same analysis for the second vendors (Siemens Healthineers) 7T human MRI system (7T Terra.X). Unlike the first vendor, the second-order shim coils from the second vendor do not align well with the ideal spherical harmonic functions. Instead, noticeable deviations, polarization swaps and field rotations are evident, indicating that the actual field patterns differ from their canonical models. This discrepancy suggests potential issues related to B_0_ shim coil design and manufacturing tolerances. Consequently, the second-order B_0_ shimming for vendor two may be less effective, as the computed B_0_ shim current solutions by the vendor integrated shim software assumes an ideal shim field basis that does not hold true considering the actual hardware. Third-order B_0_ shim coils of the second vendor, much like those from vendor one, also show strong linear cross terms. These impurities further complicate the B_0_ shimming process and limit the practical benefit that could be obtained from employing third-order B_0_ shim terms. In principle, third-order B_0_ shimming should refine the field uniformity, complementing second-order solutions by addressing subtler inhomogeneity patterns. However, if these higher-order B_0_ shim coils introduce dominant linear fields, they effectively negate the advantages that might be gained, creating a trade-off scenario where employing these coils may yield marginal net improvements or even degrade the field uniformity in certain regions.

**Figure 3.**
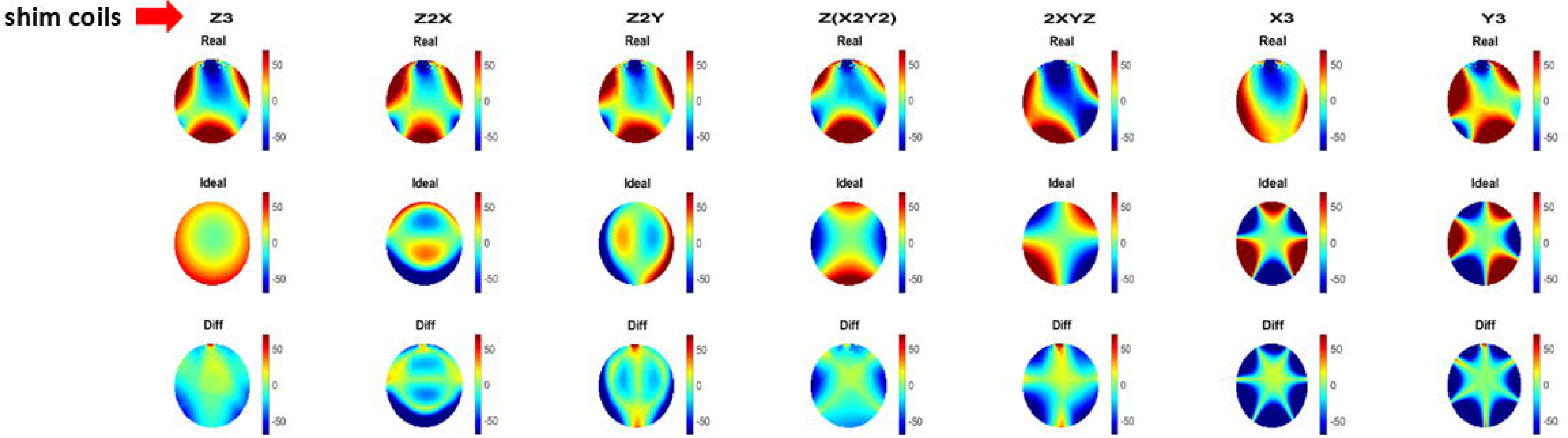
shows the real B_0_ shim fields (top row) for the third order SH B_0_ shim coils from the 7T Siemens Terra.X MRI system (vendor 2), the respective ideal SH fields (middle row) and the difference maps (bottom row). Moderate imperfections of the real shim fields including linear components are present leading to a slight difference between real and modeled ideal shim fields.

**Figure 4.**
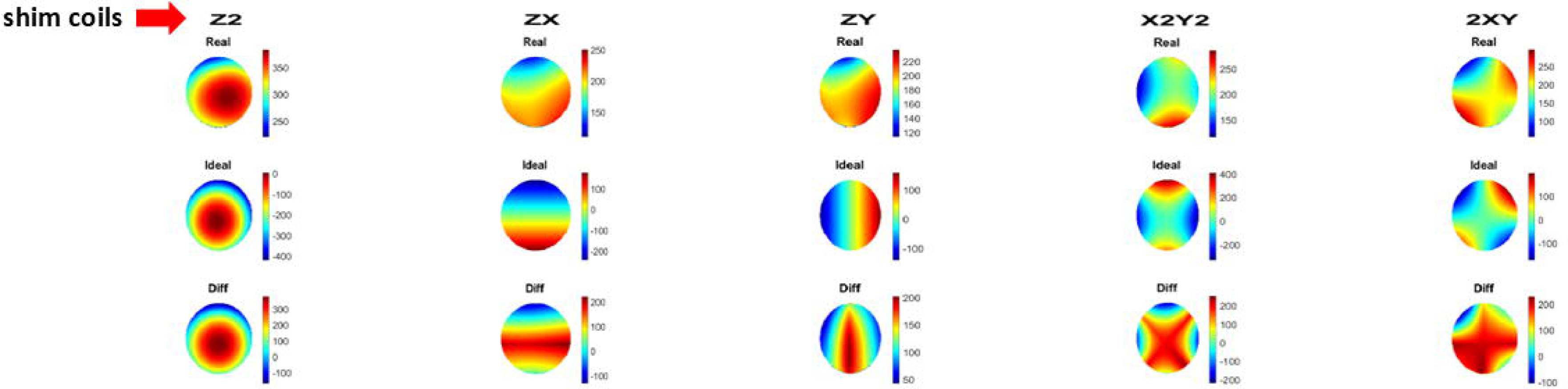
shows the real B_0_ shim fields (top row) for the third order SH B_0_ shim coils from the 7T Siemens Terra.X MRI system (vendor 2), the respective ideal SH fields (middle row) and the difference maps (bottom row). Substantial linear field components are present in the third-order real shim coils and the real third order shim fields have hardly any resemblance with the theoretical SH model

Figures 5 and 6 present a linear regression analysis comparing measured real B_0_ shim fields and ideal spherical harmonic shim fields generated by the first- and second-order shim coils from both vendors. For vendor one, most second-order B_0_ shim fields show a strong correlation with the theoretical model, although some linear gradient components are still evident. In contrast, the second-order B_0_ shim fields from vendor two display more pronounced deviations and non-uniformities. Furthermore, the data in Figure 6 highlight significant impurities within the third-order shim coils for both vendors, as demonstrated by low R² values that indicate little to no correspondence between the measured real B_0_ shim fields and their ideal spherical harmonic models.

**Figure 5.**
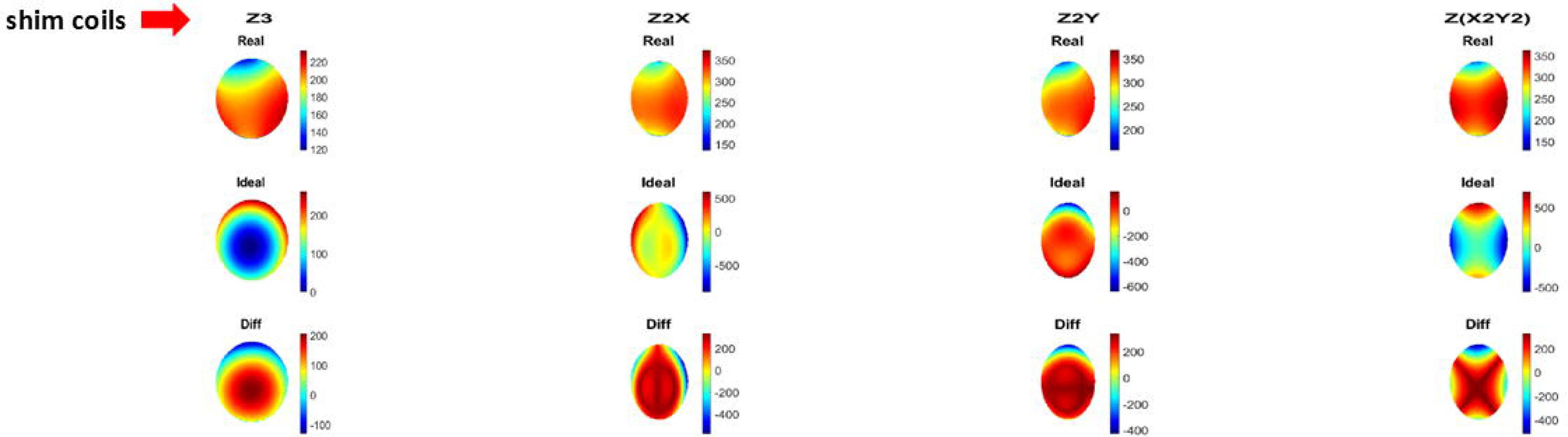
shows a linear regression analysis between measured real B_0_ shim fields and ideal B_0_ shim fields generated by first and second order B_0_ shim coils of 7T human MRI scanners from (a) vendor 1 and (b) vendor 2. Here the comparison indicates that second order shim coils from vendor 2 produce more distinct deviations of the real from the ideal shim fields than those of vendor 1.

**Figure 6.**
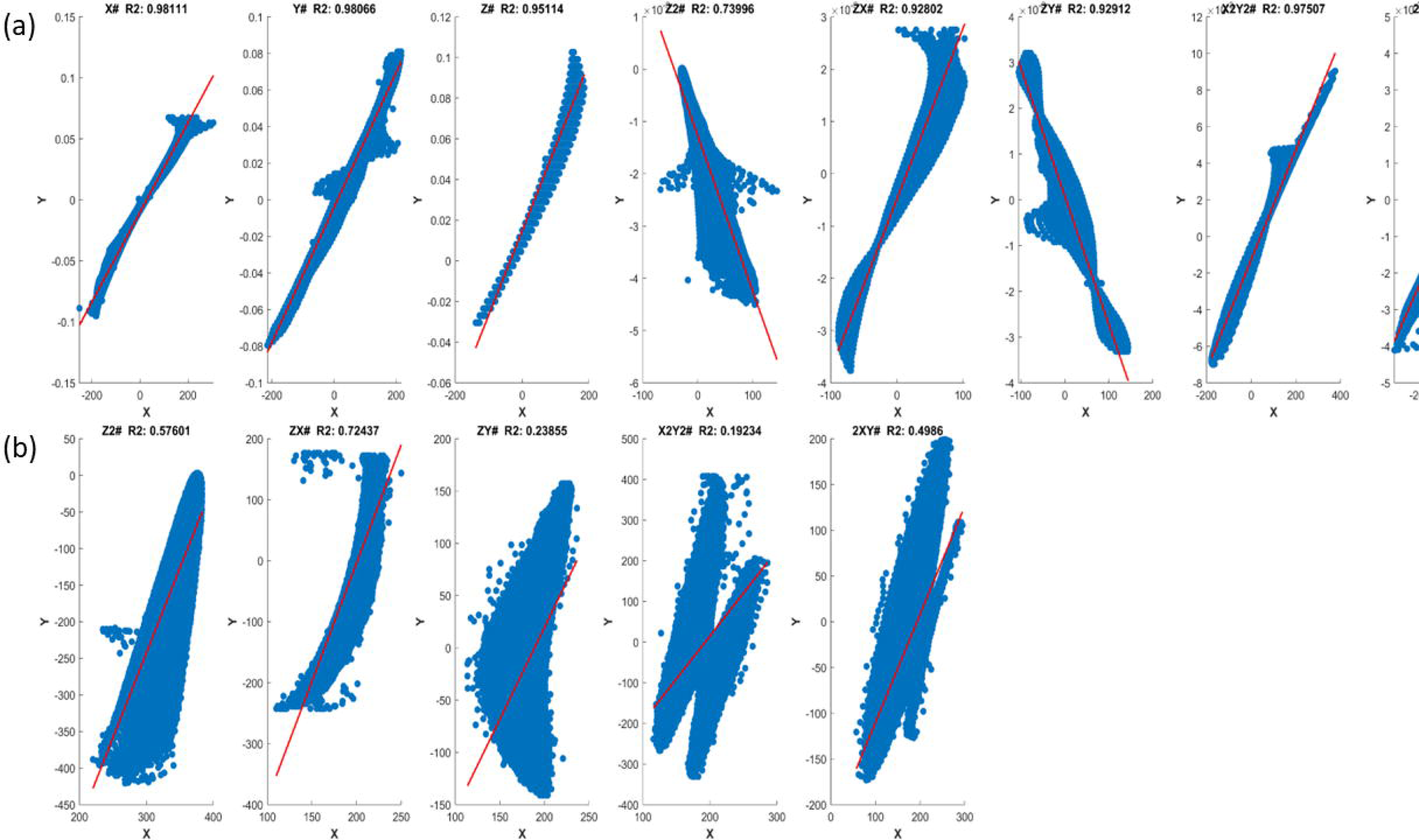
shows a linear regression analysis between measured real B_0_ shim fields and ideal B_0_ shim fields generated by third order SH shim coils of (a) vendor 1 and (b) vendor 2. There is no correlation between measured real shim fields and ideal shim field models in both vendors.

Figures 7 and 8 provide a direct comparison between the empirically measured B_0_ shim fields generated by individual B_0_ shim coils and their corresponding ideal spherical harmonic models, derived from both vendors 3T human MRI scanners. Specifically, Figure 7 compares the measured and ideal B_0_ shim field distributions for each coil in a 3T human imaging system from vendor one. Similarly, Figure 8 presents an analogous comparison for each shim coil in a 3T human imaging system from the second vendor. Both vendors 3T SH B_0_ shim systems show a better performance than their 7T counterparts with the SH in-bore B_0_ shim system of vendor one showing the highest purity.

**Figure 7.**
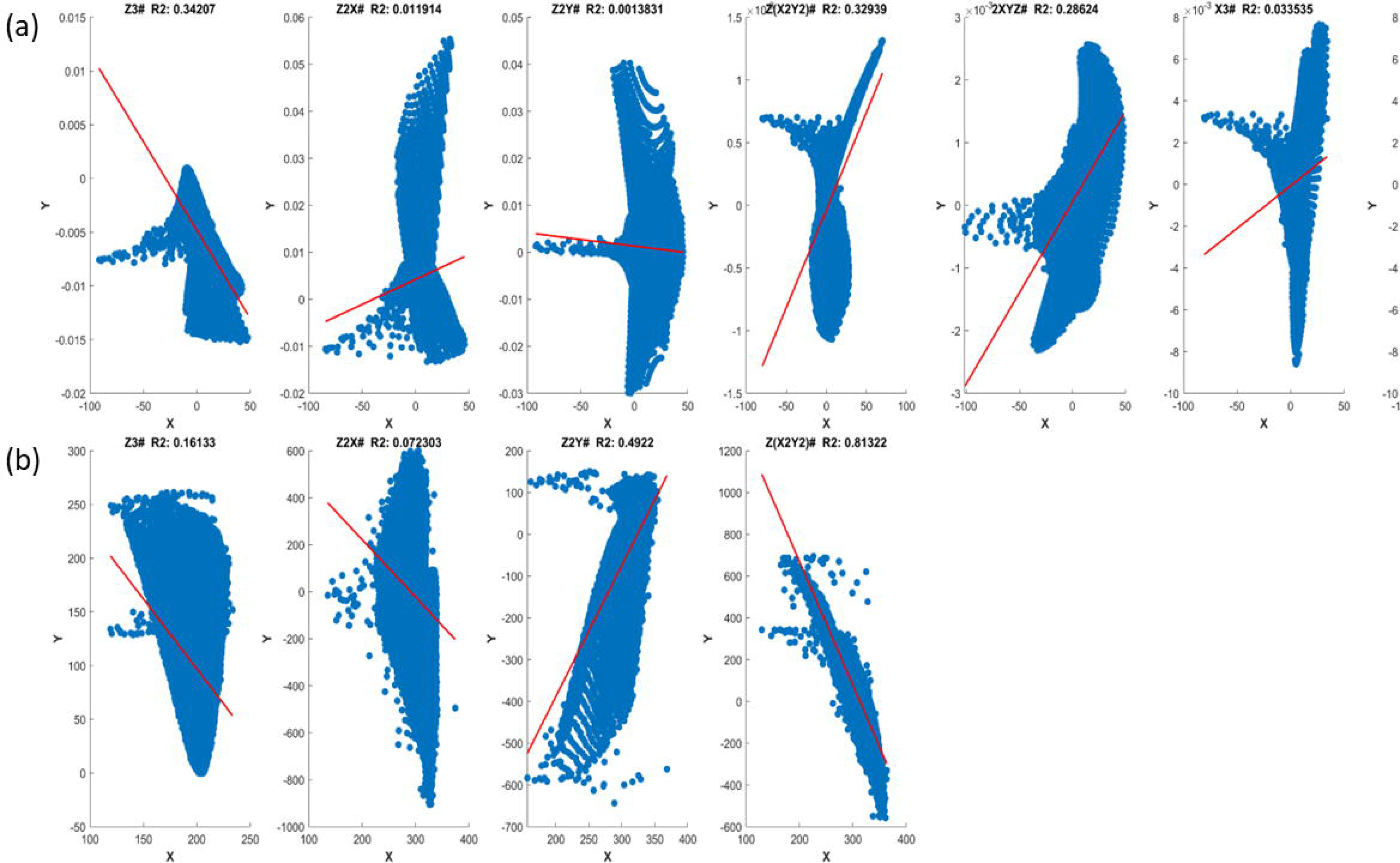
shows the real B_0_ shim fields (top row) for the first and second order SH B_0_ shim coils from the 3T Philips Ingenia MRI system (vendor 1), the respective ideal SH fields (middle row) and the difference maps (bottom row). All real shim fields almost perfectly resemble the respective ideal SH shim field models.

**Figure 8.**
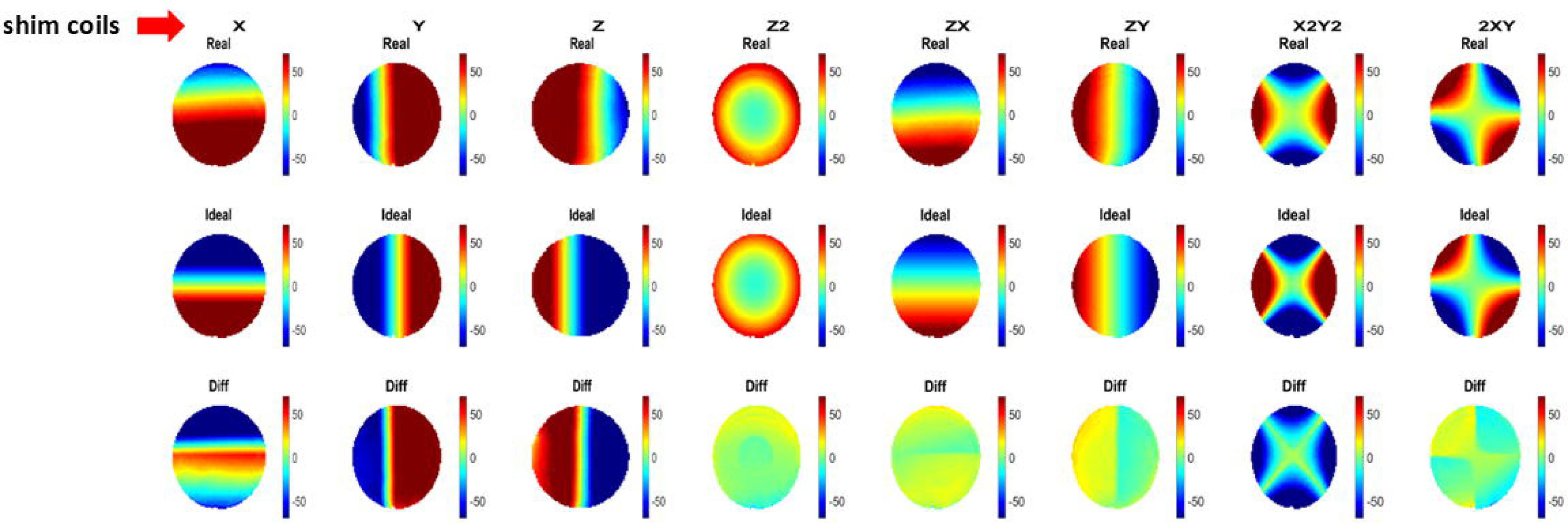
shows the real B_0_ shim fields (top row) for the first and second order SH B_0_ shim coils from the 3T Siemens Prisma MRI system (vendor 1), the respective ideal SH fields (middle row) and the difference maps (bottom row). While first order real shim fields almost perfectly resemble the respective ideal SH shim field models, real second order shim fields show moderate deviations from the ideal shim field models.

Figure 9 presents the respective linear regression analysis correlating the measured real and ideal B_0_ shim fields produced by first- and second-order coils from 3T human MRI scanners manufactured of vendor one (3T Philips Ingenia) and two (3T Siemens Prisma). In the case of vendor one, all real shim field terms closely correspond to their respective theoretical spherical harmonic models. By contrast, the shim fields from vendor two exhibit weaker associations, indicating that their measured distributions deviate more notably from the ideal shim field patterns. The correspondence between real and ideal shim fields is better for both 3T systems in comparison to the 7T human MRI scanners of the same vendor.

**Figure 9.**
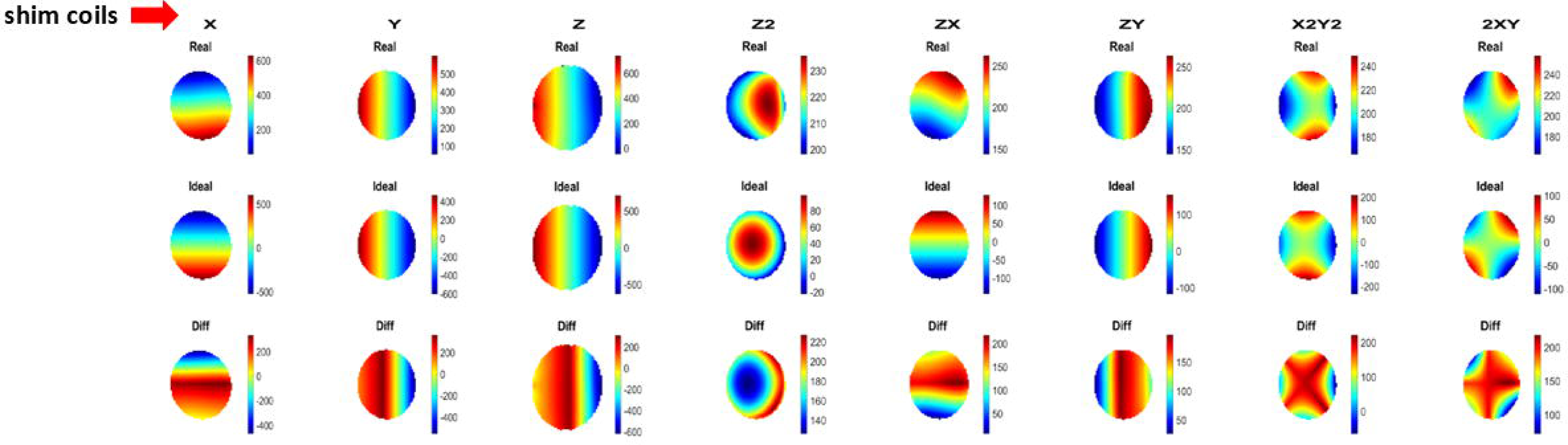
shows a linear regression analysis between measured real B_0_ shim fields and ideal B_0_ shim fields generated by first and second order B_0_ shim coils of 3T human MRI scanners from of (a) vendor 1 and (b) vendor 2. Here the comparison indicates that second order shim coils from vendor 2 produce more distinct deviations of the real from the ideal shim fields than those of vendor 1.

Supplemental Figure 5 presents the full calibration matrices for both vendors’ 7 T B_0_ shimming systems, each normalized by its coil’s self-term sensitivity. These matrices quantify the individual contributions of all shim coils to the resultant B_0_ field. In Figure 5(a), the second-order terms of vendor one align closely with their ideal spherical-harmonic patterns, confirming high spectral purity; only low-amplitude interactions between first- and second-order coils are observed, and these are too weak to perturb the overall shim solution. Examination of the third-order submatrix of the same vendor, however, reveals significant crosstalk into both linear (first order) and second-order components, indicative of geometric coupling of higher-order shim fields into lower-order ones. Figure 5(b) further highlights third-to-second-order interactions in the SH shim system of vendor 2, although incomplete calibration data prevent a full assessment of third-to-first-order coupling.

Supplemental Figure 6 shows the analogous self-term–normalized calibration matrices for vendor 1 (Fig. 6(a)) and vendor 2 (Fig. 6(b)) at 3T. In both cases, off-diagonal elements are negligible compared to the 7T systems, demonstrating minimal cross-term contamination and underscoring the superior purity and isolation of SH shim terms in these lower-field systems.

To contextualize these findings, Figure 10 illustrates B_0_ shim simulations considering ideal shim fields with shim solutions considering the real shim fields with shim solutions obtained (i) assuming ideal shim fields and (ii) assuming the real shim field calibration matrices across first, second and third-order coil configurations. Along with that, we have acquired B_0_ maps using second order shimming assuming real and ideal calibration matrix with different shim polarities. These simulations were based on shimmed B_0_ field maps of vendor one as input and considered the actual shim field constraints (Supplemental Table 1) for each of the vendors. Quantitative evaluation of spatial homogeneity via the standard deviation of B_0_ (σB_0_) reveals that: First-order shimming using real fields marginally improves homogeneity relative to idealized linear gradients, reducing standard deviation (σB_0_) and narrowing the off-resonance frequency distribution. Second-order shimming yields a marked enhancement compared to third order shimming: the inclusion of quadratic terms further contracts the frequency histogram and substantially lowers σB_0_, indicating tighter field uniformity. Third-order shimming, however, paradoxically degrades performance in vivo. Strong linear contamination arising from imperfect higher-order coil profiles elevates σB₀ and induces a systematic shift in the measured water resonance frequency (F_0_) away from predicted values. Establishing the correct polarity of the shim system is a critical factor in achieving effective B_0_ shimming, as improper polarity can introduce field distortions rather than compensating inhomogeneities.

**Figure 10.**
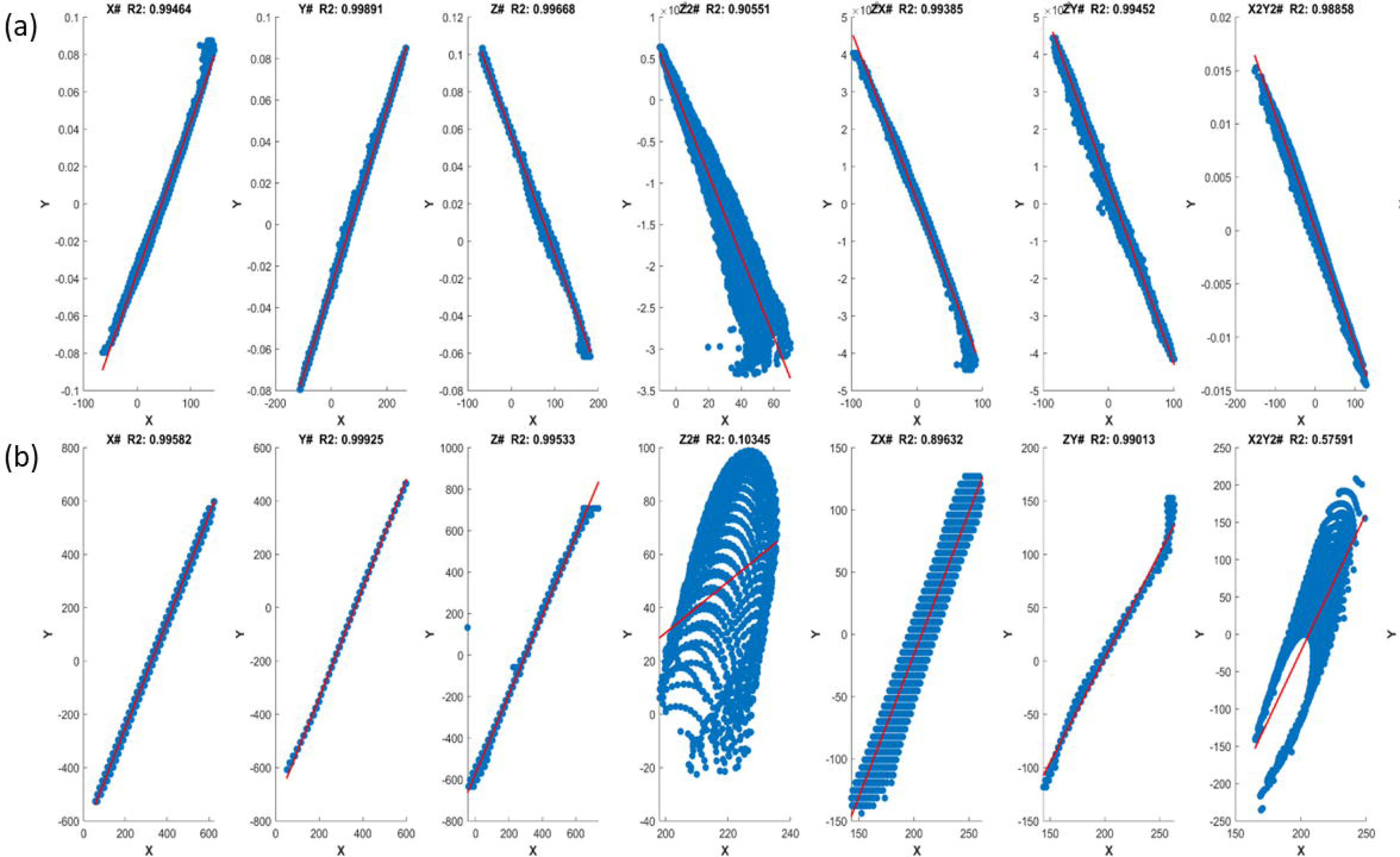
shows the comparison of B_0_ maps acquired with different shim orders acquired with real shim calibration and ideal shim calibration along with shim system polarity. The first four figures starting from left were acquired with different shim polarity and ideal and real calibration matrix. Three figures from the right were acquired using real calibration matrix and correct shim polarity with different shim orders. As we can see with using real shim calibration, homogeneity across shimmed slices is improved. Also, due to higher impurity in the higher order shim coils, we don’t see much improvement in the B_0_ maps acquired though.

These results demonstrate that, while second-order corrections confer clear gains in field homogeneity at current human 7T MRI scanners, the present very strong third-order shim coil imperfections limit incremental benefit and introduce bias in F₀ measurement and may even deteriorate the shim performance. Optimizing third-order coil purity in the shim coil design process will therefore be essential to realize true high-order B_0_ shimming gains at 7T whole-body human MRI scanners.

Quantitative analysis of the calibration data revealed a highly linear transfer function between the shim coil current and the generated ΔB₀ field strength for all vendor systems. Across the entire ±1 A dynamic range, the slope (field-per-ampere gain) remained constant to within <2 % and the coefficient of determination R2 exceeded 0.99, indicating that higher-order amplitude distortions are negligible. This near-ideal proportionality simplifies the system model to a first-order gain term, enabling straightforward prediction of shim performance from the applied current vector and obviating the need for nonlinear correction terms in routine operation. Respective results are shown in supplemental Figures 1 – 4. The maximum shim currents differ substantially between the vendors are shown in supplemental Table 1.

## Discussion

In this study, we characterized and compared the performance of second and third-order SH B_0_ shim hardware from four different commercial 7T and 3T whole-body MRI systems. The analysis revealed distinct variations in the accuracy and purity of the B_0_ shim fields generated by these systems. Specifically, for one of the evaluated vendors, the measured first- and second-order spherical harmonic shim fields closely approximated their corresponding ideal fields at 3T and 7T. In contrast, the other vendor’s second-order shim coils displayed noticeable deviations at both field strength in the form of polarization inversions and spatial rotations relative to the theoretical models. In the plotted comparison of the acquired field maps with theoretical SH models shown in Figures 3, 4 and 8 we have corrected the polarity inversions for better illustration of the actual shim field impurities. However, polarity corrections must be considered while determining shim current values^34^. For both vendors 7T shim coil designs for second order spherical harmonics show higher shim field impurities than respective 3T designs.

A more pervasive challenge emerged for third-order B_0_ shimming, where both vendors’ B_0_ shim coils introduced substantial linear field components. These unintended linear contributions not only compromised the expected enhancement in B_0_ homogeneity but also introduced significant zero-order frequency shifts. Such alterations impede the intended improvements in image quality, as they effectively counteract the benefits gained from lower-order B_0_ shimming and can even degrade overall B_0_ field uniformity.

The findings underscore the need for robust real shim field calibration protocols especially at 7T human MRI systems. While second-order image-based B_0_ shimming can achieve meaningful improvements if the actual shim fields are thoroughly mapped and modeled, this step currently requires the use of custom or third-party B_0_ shimming software, as vendor-integrated methods do not offer direct real field calibration options. Without proper calibration, even second-order B_0_ shimming may fail to meet its full potential when using current 7T SH shim coil designs (Figure 10). In practical terms, these results underscore the importance of conducting a comprehensive calibration of B_0_ shim coils and use of their real shim fields in shim algorithms rather than relying solely on ideal spherical harmonic assumptions upto his. In case of high purity 2^nd^ order SH shim coils only available at 3T cross-terms may induce only negligible adjustments to the calculated shim currents implying that an explicit calibration matrix may be unnecessary in such systems ^20^. With calibration we can compensate for smaller deviations of the real from ideal B_0_ shim fields causing geometric coupling between B_0_ shim coils. However, if the deviations from the intended shim field pattern is very strong as the herein reported interaction between third order shim coils with first order and second shim coils as shown in the supplement figure 5 (a) and (b) calibration cannot counter that. It is hence preferred to have higher order B_0_ shim coils designed with purer shim fields resembling spherical harmonics terms.

More critically, the results caution against employing third-order B_0_ shimming at the current 7T whole-body human MRI scanners using vendor-provided hardware configurations in their current form. The substantial discrepancies between measured real and ideal third-order B_0_ shim fields, coupled with the pronounced linear components, mean that attempts to refine the field with third-order terms may actually deteriorate rather than improve the achieved B_0_ homogeneity.

Our findings also refine the current understanding of interactions between gradient coils and different shim coils that were previously characterized with dynamic field-monitoring techniques^19,29,27,33,34^. By acquiring the complete steady-state calibration matrix, we confirm the pronounced linear cross-terms between third order and linear terms reported earlier^19,20,36^ and demonstrate that they predominantly originate from geometric impurities instead of inductive coupling an ambiguity that earlier studies using spatio-temporal field monitoring could not resolve. The development of more sophisticated B_0_ shim coil designs could mitigate linear impurities, ensuring that higher-order B_0_ shim coils fulfill their intended role in fine-tuning the magnetic field ^21, 26^ . Future design methods must focus on improving the purity of higher-order B_0_ shim fields to fully capitalize on their theoretical advantages, thereby enabling ultra-high-field MRI scanners to deliver consistent, high-quality images and accurate spectroscopic data.

From an engineering and calibration standpoint, the finding of a linear relationship between the input current to the B_0_ shim coils and their corresponding B_0_ shim field strengths is a welcome characteristic. It simplifies the process of acquiring the real shim field data and modeling the shim coil responses, as one does not need to account for nonlinear scaling factors, at least within the current amplitude range tested. Nevertheless, linearity in amplitude does not solve the issue of spatial field distortions due to inaccurate harmonic patterns.

In forthcoming studies, we will rigorously quantify the degree to which third-order shim coils couple into the linear B₀ field terms by computing coupling coefficients for each third-order basis function. We will then assess what fraction of these cross-term interactions expressed as a percentage of the self-term amplitude—can be effectively compensated by the full calibration matrix. Finally, by analyzing the residual field error as a function of coil order and coupling magnitude, we will pinpoint the threshold beyond calibration of higher-order shim coils yields negligible improvement in prescribed shim currents, thereby defining the point of diminishing returns for third-order shim coils.

## Conclusion

In summary, accurate calibration of actual B_0_ shim fields is strongly advised for successful higher-order B_0_ shimming in commercial 3T and 7T human whole-body MRI systems. However, higher order B_0_ shim coils containing substantial shim field impurities including linear gradients such as the investigated third order SH shim coils reported here will limit the positive impact of real field shim calibration matrix. In these cases, a substantial improvement of the design and implementation of third-order shim coils at 7T human MRI scanners is needed to fully realize the theoretical advantages of higher-order B_0_ shimming. Enhancements in the purity of future generations of commercial B_0_ shim systems at 3T and 7T will ultimately translate into superior field homogeneity, reduced imaging artifacts, and improved overall scan quality.

## Supplemental material

### Table Captions

Supplement Table 1 shows shim coil sensitivity and maximum shim field strength of all investigated 3T and 7T spherical harmonic B_0_ shim system.

Supplemental Figure 1: Linear relationship between applied shim current and generated shim field strength from 7T Philips dSync MRI system. The percentage change in theoretical and measured current is less than 2% and the shim system shows a highly linear performance.

Supplemental Figure 2: Linear relationship between applied shim current and generated shim field from 7T Siemens Terra.X MRI system. The percentage change between theoretical and measured current is less than 2% and the shim system shows a highly linear performance.

Supplemental Figure 3: Linear relationship between applied shim current and generated shim field from 3T Philips Ingenia MRI system. The percentage change between theoretical and measured current is less than 2% and the shim system shows a highly linear performance.

Supplemental Figure 4: Linear relationship between applied shim current and generated shim field from 3T Siemens Prisma MRI system. The percentage change between theoretical and measured current is less than 2% and the shim system shows a highly linear performance.

Supplemental Figure 5: Shim calibration matrix for (a) 7T Philips dSync and (b) 7T Siemens Terra X respectively.

Supplemental Figure 6: Shim calibration matrix for (a) 3T Philips Ingenia and (b) 3T Siemens Prisma MRI system respectively.

Supplemental Figure 7: comparison of B_0_ maps acquired through different shimming with ideal and real shim calibration matrix, considering with/without correct polarity of the shim system. The B_0_ maps were acquired though slices above corpus collosum. We can see B_0_ maps acquired after considering real calibration matrix and with correct polarity of shim system are more homogenous.

