## Supplemental material for "Characterization of real B_0_ shim fields ^generated^ by higher order B_0_ shim systems of whole body human 3T and 7T MRI systems"

**Table Captions:**

Supplement Table 1 shows shim coil sensitivity and maximum shim field strength of all investigated 3T and 7T spherical harmonic $B_0$ shim system.

Commented [AH1]: I think we need this for the analysis to be shown in Figure 10.

Commented [MJ2R1]: Ivan told me once, you can not add vendors specification into paper. This paper has maximum shim capacity and sensitivity.

Supplemental Figure 1: Linear relationship between applied shim current and generated shim field strength from 7T Philips dSync MRI system. The percentage change in theoretical and measured current is less than 2% and the shim system shows a highly linear performance.

Supplemental Figure 2: Linear relationship between applied shim current and generated shim field from 7T Siemens Terra.X MRI system. The percentage change between theoretical and measured current is less than 2% and the shim system shows a highly linear performance.

Supplemental Figure 3: Linear relationship between applied shim current and generated shim field from 3T Philips Ingenia MRI system. The percentage change between theoretical and measured current is less than 2% and the shim system shows a highly linear performance.

Supplemental Figure 4: Linear relationship between applied shim current and generated shim field from 3T Siemens Prisma MRI system. The percentage change between theoretical and measured current is less than 2% and the shim system shows a highly linear performance.

Supplemental Figure 5: Shim calibration matrix for (a) 7T Philips dSync and (b) 7T Siemens Terra X respectively.

Supplemental Figure 6: Shim calibration matrix for (a) 3T Philips Ingenia and (b) 3T Siemens Prisma MRI system respectively.

Commented [AH3]: Add (a) and (ab) into the Figures.

Commented [MJ4R3]: What?

Commented [MJ5R3]: It is there

Commented [AH6]: Add (a) and (ab) into the Figures.

Commented [MJ7R6]: What?

Commented [MJ8R6]: It is there

Supplemental Figure 7: comparison of $B_0$ maps acquired through different shimming with ideal and real shim calibration matrix, considering with/without correct polarity of the shim system. The $B_0$ maps were acquired though slices above corpus collosum. We can see $B_0$ maps acquired after considering real calibration matrix and with correct polarity of shim system are more homogenous.

### Supplement Table 1

Table 1

| Vendors | Philips | | | | Siemens | | | |
|---|---|---|---|---|---|---|---|---|
| **Field Strength** | **7T** | | **3T (Ingenia)** | | **7T (terra X)** | | **3T (Prisma)** | |
| **Shimcoils** | coil sensitivity (A*m^n/mT) | field strength(max) mT/m^n | coil sensitivity (A*m^n/mT) | field strength (max) mT/m^n | coil sensitivity (A*m^n/μT) | field strength (max) μT/m^n | coil sensitivity (A*m^n/μT) | field strength (max) μT/m^n |
| X | 15.2864 | ±1 | 18.6109 | ±1 | | | 6.3 | ±2300 |
| Y | 15.3392 | ±1 | 18.9332 | ±1 | | | 6.28 | ±2300 |
| Z | 15.4068 | ±1 | 19.2432 | ±1 | | | 6.1 | ±2300 |
| Z2 | -2.1916 | ±4.5 | -2.3069 | ±2.1 | 7.48 | | 2.014 | ±4959 |
| ZX | 1.3342 | ±7.5 | 0.9335 | ±5.35 | 3.74 | | 2.814 | ±3551 |
| ZY | -1.3423 | ±7.5 | 0.9496 | ±5.25 | 3.69 | | 2.85 | ±3503 |
| X2Y2 | 1.3264 | ±7.5 | -2.2739 | ±2.15 | 3.69 | | 2.814 | ±3551 |
| 2XY | 1.3605 | ±7.3 | -2.2492 | ±2.20 | 3.64 | | 2.864 | ±3487 |
| Z3 | -3.3434 | ±2.96 | | | 15.23 | | | |
| Z2X | -2.7405 | ±3.6 | | | 14.01 | | | |
| Z2Y | -2.7675 | ±3.6 | | | 14.01 | | | |
| Z(X2Y2) | 0.438 | ±22.8 | | | 14.01 | | | |
| 2XYZ | 0.4405 | ±22.7 | | | | | | |
| X3 | 0.9617 | ±10 | | | | | | |
| Y3 | 0.9924 | ±10 | | | | | | |

### Supplemental Figure 1

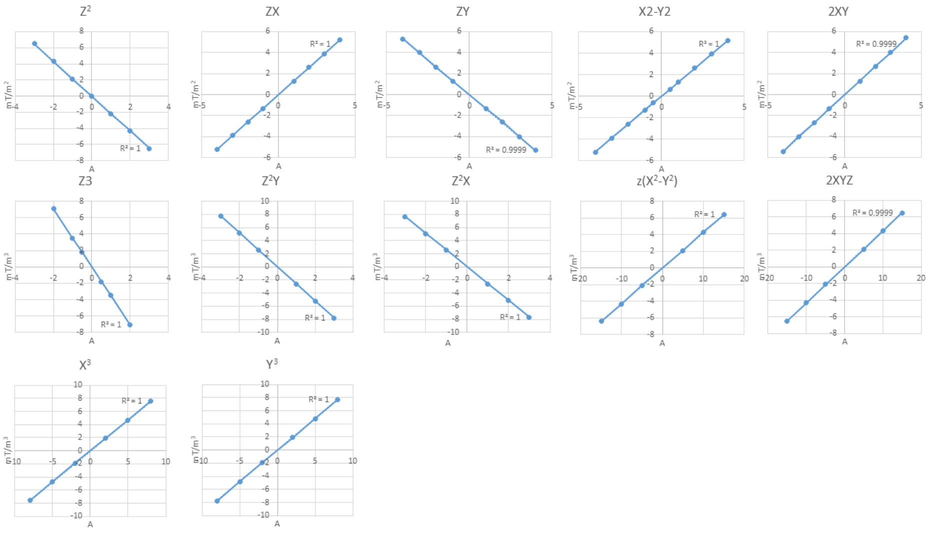

Supplemental Figure 2

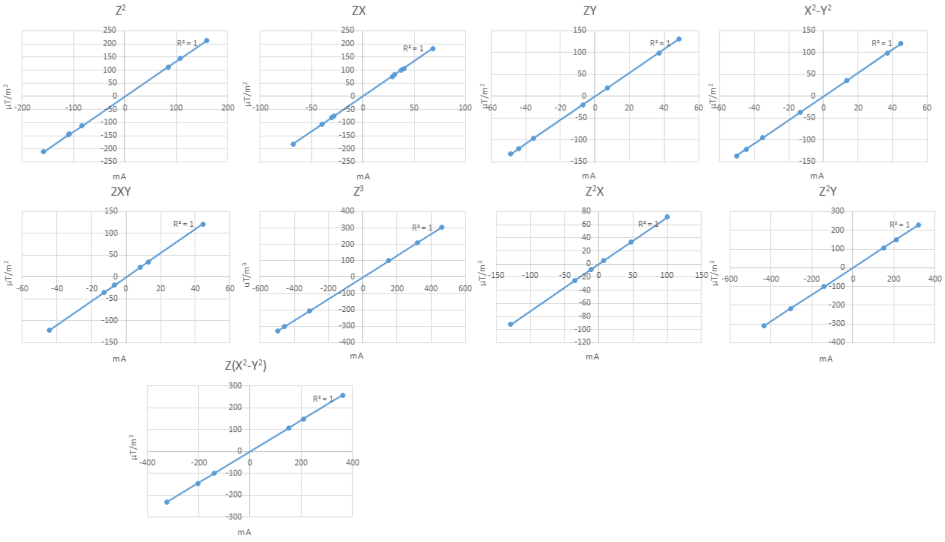


Supplemental Figure 3

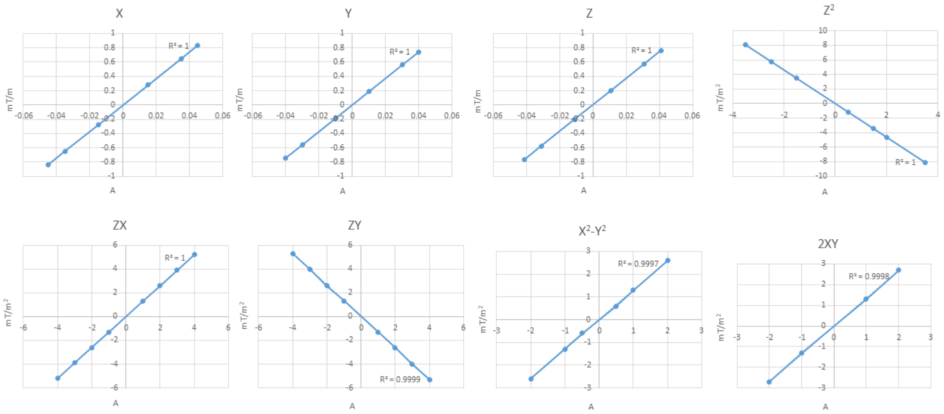


Supplemental Figure 4

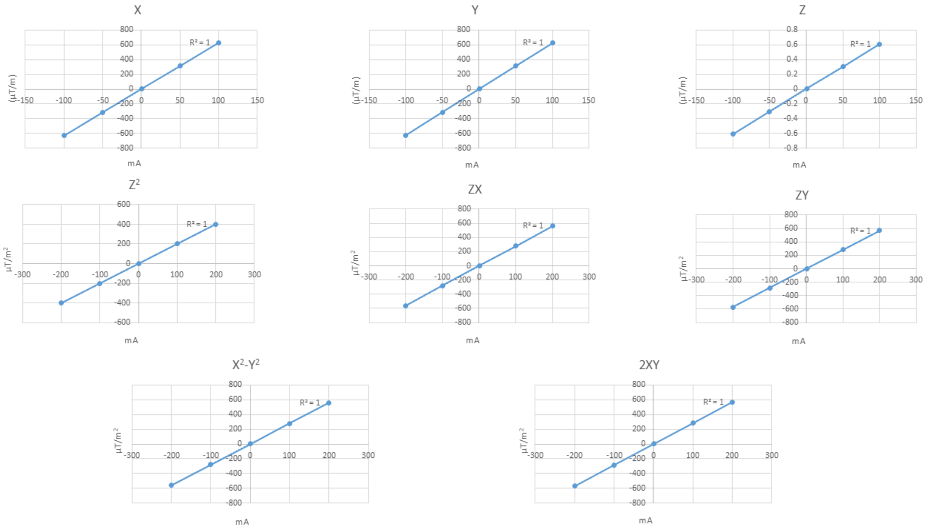

### Supplemental Figure 4

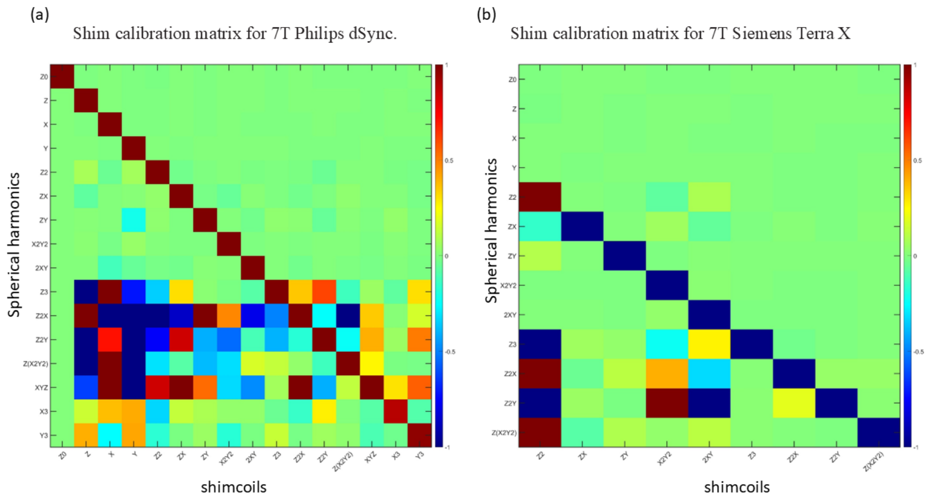


### Supplemental Figure 5

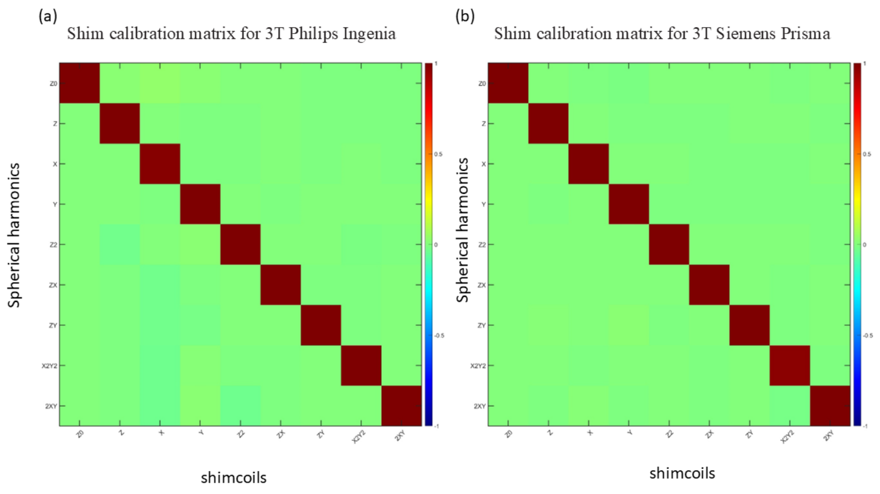

Supplemental Figure 6

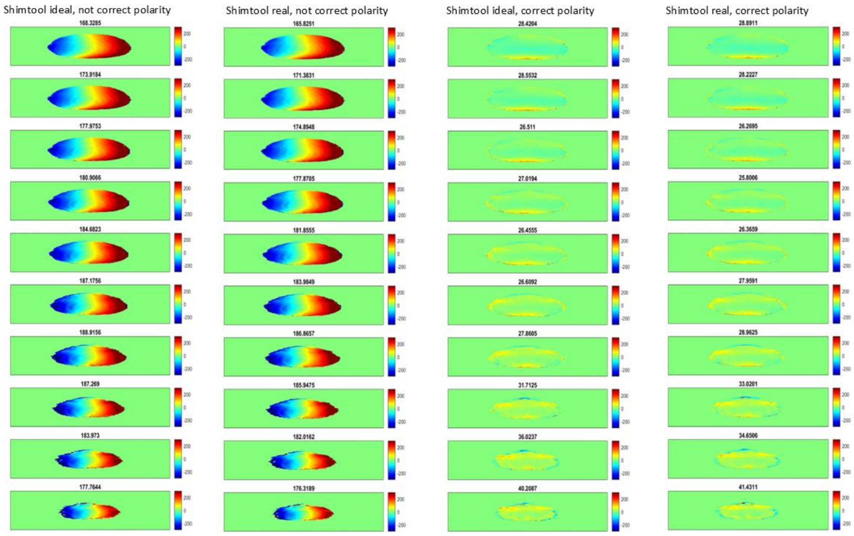
